# Auditory Brainstem Development in Autism: From Childhood Hypo-Responsivity to Adult Hyper-Reactivity

**DOI:** 10.1101/2025.04.22.650041

**Authors:** Ala A. Seif, Renee Leow-Guerville, Mohammed S. Rajab, Cassandra Marceau-Linhares, Kristina Schaaf, Samantha E. Schulz, Susanne Schmid, Ryan A. Stevenson

## Abstract

**Background:** Autism Spectrum Disorder (ASD) is characterized by sensory disruptions, including auditory processing differences, which can significantly impact social, emotional, and cognitive development. This study investigates auditory brainstem development in Autistic children and adults using auditory brainstem responses (ABRs) and acoustic startle responses (ASRs), two key measures of auditory processing. We hypothesize that early hypo-responsivity in children, measured with ABRs, may lead to compensatory neural adaptations, resulting in hyper-reactivity in adulthood, measured by ASRs.

**Methods:** The study included 40 Autistic children, 57 non-Autistic children, 20 Autistic adults, and 21 non-Autistic adults. Participants underwent peripheral hearing screening, ABR testing at slow and fast click-rates, and ASR measurements. ABR wave and ASR amplitude were analyzed. Statistical analyses included mixed-model ANOVAs and Spearman’s correlations to examine group differences and associations with age.

**Results:** Autistic participants showed trending differences in auditory brainstem timing relative to non-autistic participants, with no differences in brainstem amplitude at any wave or rate. *Post hoc* slope comparisons revealed that non-Autistic participants showed positive developmental trajectories in Wave I latency at both rates, while Autistic participants showed flat trajectories, a pattern directionally consistent with developmental convergence. A sex-specific group difference was observed at Wave I slow rate, with Autistic females showing longer latencies than non-Autistic females (d = 1.69), with no difference in males. In contrast, ASR amplitude showed diverging developmental trajectories between groups, with Autistic participants showing a positive age-related increase and non-Autistic participants showing a negative age-related decrease (slope contrast: t(99) = 2.16, p = .033, η^2^p = .05), consistent with a group difference emerging in adulthood. ABR amplitude was positively correlated with ASR amplitude in non-Autistic participants (Wave I: rho = +0.30; Wave V: rho = +0.25) but negatively correlated in Autistic participants (Wave I: rho = −0.19; Wave V: rho = −0.20), with the between-group difference in correlation direction surviving FDR correction for both waves (Fisher’s z p_adj_ = .035).

**Conclusion:** The findings are consistent with the hypothesis that Autistic children experience auditory brainstem hypo-responsivity, which may normalize in adulthood but lead to maladaptive hyper-reactivity. These results highlight the role of early auditory disruptions in shaping long-term sensory processing and reactivity in Autism, emphasizing the need for further research into the neural mechanisms underlying these differences.

## Introduction

Autism is a neurodevelopmental condition characterised by challenges in social interaction, communication, and the presence of restricted and/or repetitive behaviors and interests. Sensory disruptions, such as hyper- and hyposensitivities, were only included in the most recent version of the diagnostic criteria under the umbrella of restricted and repetitive behaviors^1,2^. Sensory disruptions are well-documented in clinical^3,4^ and experimental measures^3,5^. Moreover, difficulties in sensory processing can cascade to affect social communication and interactions^6–9^, education^10–13^, and everyday activities^10,14^. Moreover, they are associated with higher rates of social-emotional dysregulation^15,16^, anxiety^16–18^, and other mental health difficulties^19,20^. Sensory disruptions are often categorized into three subcategories, hyperresponsiveness, hyporesponsiveness, and sensory seeking^19^. They are reported in multiple modalities^21^ such as olfactory^22,23^, tactile^24–26^, auditory^27^, visual^28^, as well as in the interplay of those modalities^29–34^.

Auditory processing plays a crucial role in cognitive development^35^, impacting how individuals interpret, understand, and respond to sound stimuli in their environment. Disruptions in auditory processing during early development and critical periods is especially impactful^36–38^. These periods close after structural consolidation, reducing future plasticity as the brain reaches adulthood. Although critical periods provide an exceptional time window for learning and consolidation, they also represent a period of great vulnerability for the developing brain^39^. This study focused on the auditory sensory disruptions at the level of the brainstem across development. There are multiple studies reporting alterations in the ascending auditory brainstems in Autistic individuals, including slower neurotransmission^40–43^ and atypical cell structures^44–46^. In addition, there is evidence from animal models of Autism indicating reduced responsivity of the ascending auditory pathway^47,48^.

The functionality of the auditory brainstem pathway is commonly measured through the auditory brainstem response (ABR) a well-characterized auditory evoked potential of the brainstem. The ABR output is a waveform with five major wave peaks that have well-established bio generators. Wave I is linked to the distal portion of the auditory nerve with an absolute latency of 1.5 msec^49,50^. While wave II is associated with the proximal portion of the auditory nerve and cochlear nucleus with an absolute latency of 2.5 msec^50,51^. Wave III is generated by the cochlear nucleus and superior olivary complex with an absolute latency of 3.5 msec^52^. Wave IV is associated with the superior olivary complex and ascending auditory fibers of the lateral lemniscus with an absolute latency of 4.5 msec^52^. Wave V corresponds to the lateral lemniscus and inferior colliculus with an absolute latency of 5.5 msec ^53–55^ (Figure 1). The latency of the wave is indicative of the speed of neurotransmission while the amplitude of the waves is indictive of neural responsivity^53^.

**Figure 1:**
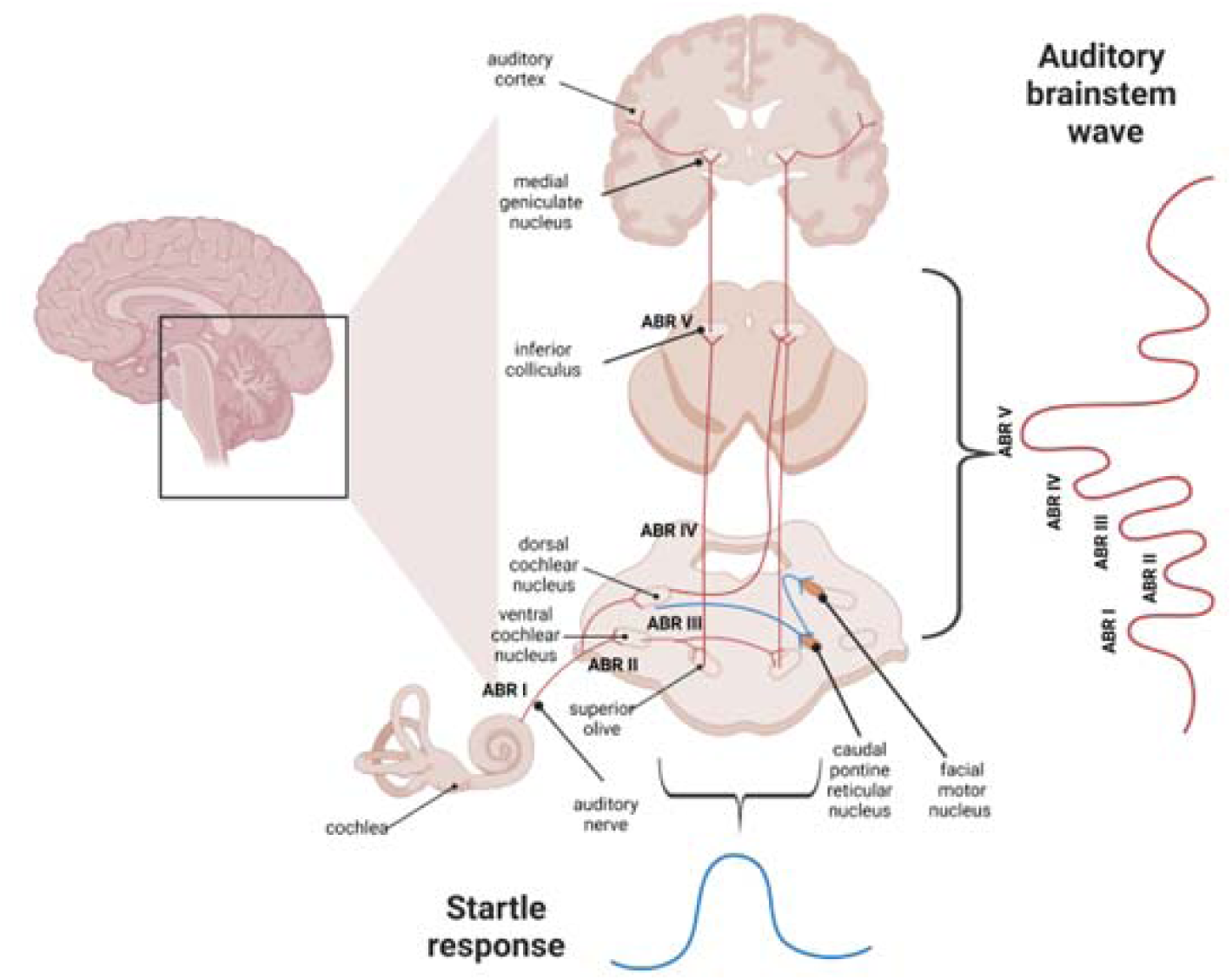
The pathways involved with the acoustic startle reflex and the auditory brainstem response.

ABRs are measured as a common newborn hearing test to provide information about brainstem maturation. Given this, is has been possible to detect differences in ABRs in infants later diagnosed with Autism^56–58^. The ABR reports in Autism, however, have been inconsistent, with some reporting no group differences and others reporting delayed and/or muted ABR waves for Autistic individuals^59^. The inconsistency could be owed to the heterogeneity of Autism, the variation in the ABR parameters, or the participant ages in the studies. Studies that looked at ABRs in younger samples such as newborns^56–58^, toddlers^60,61^, or school-aged^62,63^ children report increased ABR peak latencies. However adult study results are less consistent^64–66^. This could be due to a delay in the development of the ascending auditory pathway.

There is also evidence supporting increased behavioral responses to sound associated with Autism, including brainstem-mediated reflexive responses. These responses are commonly measured by the acoustic startle response (ASR), which is an evolutionary conserved protective response to sudden auditory stimuli^4,67–69^. The primary startle pathway includes spiral ganglion cells innervating the cochlea and synapsing on secondary auditory neurons in the cochlear nucleus which innervate the ventrolateral caudal pontine reticular nucleus (PnC)^70^. PnC giant neurons project to the spinal cord where they ipsilaterally innervate motor neurons^70^ (Figure 1). The startle response is a measure of acoustic reactivity and is typically quantified in humans using electromyography (EMG) of the eyeblink, in which electrodes measure orbicularis oculi muscle activity. Autistic individuals are reported to have increased startle reactivity. According to the literature, Autistic individuals have been reported to express acoustic hyperreactivity^67^ and hyposensitivity to complex stimuli in the ascending auditory pathway^71^, measured by the auditory brainstem response. It is unclear if these two seemingly contradicting constructs are explained by the heterogeneity of Autism.

This study focused on the auditory brainstem development by investigating the auditory brainstem response and the acoustic startle response in a sample of Autistic children and Autistic adults. According to previous report, we expect that Autistic children will have hyporesponsivity to sounds quantified by muted and delayed ABR wave peaks compared to non-Autistic children, while Autistic adults will have normalized ABR. We hypothesize that this hypo-responsivity in Autistic children will lead to the development of an adaptive compensatory gain of auditory signals downstream of the ABR. As the hyporesponsivity of the ABR gets resolved with age, the adaptive compensatory gain would become maladaptive. Therefore, we predict that the Autistic adults will have increased behavioral reactivity measured by startle response compared to non-Autistic adults. Finally, we hypothesize that ABR parameters will negatively correlate with age while behavioural reactivity will positively correlate with age for the Autistic participants.

## Methodology

### Participants

This study includes 40 Autistic children and 57 non-Autistic children. It also included 20 Autistic adults and 21 non-Autistic adults (Table 1). All participants had normal hearing (confirmed in the study, details below) and absence of neurological disorders, history of seizures, or past head injury. Participants’ cognitive ability (IQ > 70) was assessed by the Wechsler Abbreviated Scales of Intelligence, second edition^72^ or Kaufman Brief Intelligence Test, Second Edition^73^. The study was a fully passive study therefore allowing us to include 3 Autistic participants that could not sit through the IQ test due to communication difficulties. Participants in the Autism group had a clinical diagnosis of Autism by a professional including medical doctors such as family physicians, pediatricians and psychiatrists, psychologists and psychological associates, or nurse practitioners. The diagnosis was confirmed by an Autism Diagnostic Observation Schedule, Second Edition (ADOS-2)^74^ conducted by a research-reliable administer, or a clinically reliable administer supervised by a research-reliable administer. Participants in the non-Autism group had no family history of Autism in first-degree relatives. An additional 2 participants were excluded due to ADOS-2 not confirming the Autism diagnosis. Final groups did not differ on hearing screening. Participants aged eighteen or more or the parents of younger participants gave informed assent, and children gave assent.

**Table 1:**
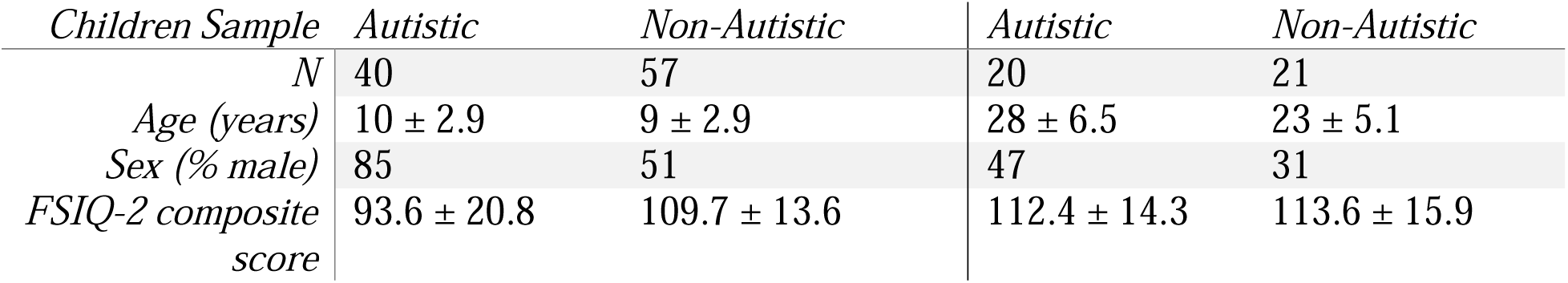
Demographic information for the sample. A subset of the adult non-Autistic sample were undergraduate university students, and they did not complete a cognitive assessment as it was assumed to be IQ >70.

| *Children Sample* | *Autistic* | *Non-Autistic* | *Autistic* | *Non-Autistic* |
| --- | --- | --- | --- | --- |
| *N* | 40 | 57 | 20 | 21 |
| *Age (years)* | $10 \pm 2.9$ | $9 \pm 2.9$ | $28 \pm 6.5$ | $23 \pm 5.1$ |
| *Sex (% male)* | 85 | 51 | 47 | 31 |
| *FSIQ-2 composite score* | $93.6 \pm 20.8$ | $109.7 \pm 13.6$ | $112.4 \pm 14.3$ | $113.6 \pm 15.9$ |

### Periphery Hearing Screening

A periphery hearing screening was conducted before being included in the study. The screening consists of a visual otoscope inspection, an audiometric assessment (Midimate 602), and a tympanometry test (MT10 by interacoustics). The evaluation criteria are (1) no perforation in the eardrum, (2) pass an audiometer screen for frequencies (250 Hz, 500 Hz, 1000 Hz, 2000 Hz, 4000 Hz, 8000 Hz) < 25 dB, and (3) have a detected tympanogram peak with compliance(ml) = 0.30-1.5, pressure (dapa) = -/+100 and ear canal volume (ECV) = 0.5-1.35. All assessments were completed for both ears and the participants were included if they passed 2 out of the 3 assessments for every ear. There were 10 participants excluded from the study based on their periphery hearing screening. They are not included in the study’s total number of participants.

### Auditory Brainstem Response

The ABR was measured using ABR Integrity™ V500 System. The acoustic stimuli were presented via ER-2 insert earphones. Presentations were made monoaurally and click pulses were repeated until a total of 2000 accepted evoked potential sweeps had been collected and accepted. The acoustic stimuli were presented at a slow click-rate (19.1 clicks/sec) and a fast click-rate (59.1 clicks/sec) at sound pressure level of 80 dB HL. Two trials were recorded for every condition from each ear. The peaks were marked by two trained personnels. The ABR variables were averaged across both trials and ears for every participant.

### Acoustic Startle Response

The acoustic startle response was recorded using SR-HLAB™ EMG. All auditory stimuli and background white noise were delivered binaurally to participants through stereophonic headphones. Startle eyeblink electromyographic responses were recorded from the left orbicularis oculi muscle. The eyeblink magnitude of every startle response was defined as the voltage of the peak electromyographic activity within a latency window of 20 to 160 msec following startle-eliciting stimulus onset. The startle onset was defined as the point of positive inflection of the volts before startle peak and the startle latency is the latency of the peak voltage. The peaks were marked automatically by the software. The paradigm consisted of a 5-minute acclimation period and consistent 60 dB white noise background. The acoustic reactivity trials consisted of white noise pulse ranging from 65 dB to 105 dB for 40 msec with 10 dB increments and all trial conditions were repeated 10 time. All trials were presented in a fixed pseudorandom order, separated by intertrial intervals of 10 to 20 sec (15 sec on average). Startle magnitude was averaged for every condition.

### Statistical Analysis

The main objective of the study was to investigate functional and behavioral measures of the auditory brainstem across development in Autism. ABR features (latency, interpeak intervals, and amplitude for Waves I, III, and V) and ASR amplitude were analysed using linear mixed models, with degrees of freedom estimated using the Kenward-Roger method.

For ABRs, each feature was modelled with between-subject predictors of diagnostic group (Autistic, non-Autistic), age, and sex, and a within-subjects factor of click rate (slow: 19.1/s; fast: 59.1/s) modelled as a repeated measure via a random intercept for participant. All two-, three-, and four-way interactions between predictors were included. Normality of residuals was assessed via Q-Q plots and the Shapiro-Wilk test; homoscedasticity was assessed via residuals versus fitted plots. Where parametric assumptions were not met, a rank transformation was applied prior to modelling.

For ASR, amplitude was log-transformed based on the same assumption-checking pipeline. A full polynomial model was run with dB level modelled as a quadric polynomial (poly (dB, 2)), sex, and age as a continuous variable.

Multiple comparisons were controlled using the Benjamini-Hochberg false discovery rate (FDR) procedure (Q = 0.05), applied within each feature family (latency, interpeak intervals, amplitude) for ABR. Trending effects were investigated via *post hoc* slope comparisons or simple effects and are treated as exploratory.

To assess the relationship between ABR amplitude and ASR amplitude, Spearman correlations were computed between rate-collapsed ABR amplitude composites and ASR amplitude at 105 dB, separately for Autistic and Non-Autistic participants. Between-group differences in correlation magnitude were tested using Fisher’s z transformation. All analyses were conducted in R; the study was pre-registered on the Open Science Framework (osf.io/hg6b2).

## Results

### ABR: Absolute and Interpeak latencies

We analyzed absolute latency of ABR (Table 2, Figure 2) wave I, wave III, and wave V and interpeak latencies (IPLs; Figure 4) between waves I and III (I–III IPL) as well as waves III and V (III-V IPL) and overall waveform of wave I and V (I-V IPL; sex × click-rate × diagnostic group × Age).

**Table 2.** ABR Latency and Interpeak Intervals — Group Means (SD)

| Feature | Rate | Autistic |  | Non-Autistic |  |
| --- | --- | --- | --- | --- | --- |
|  |  | Children | Adults | Children | Adults |
| **Wave I** | *Slow* | 1.38 (0.14) | 1.42 (0.16) | 1.34 (0.17) | 1.40 (0.11) |
|  | *Fast* | 1.41 (0.17) | 1.43 (0.26) | 1.39 (0.19) | 1.44 (0.15) |
| **Wave I'** | *Slow* | 2.03 (0.12) | 2.06 (0.19) | 1.97 (0.18) | 2.06 (0.12) |
|  | *Fast* | 2.06 (0.18) | 2.04 (0.30) | 2.01 (0.24) | 2.11 (0.14) |
| **Wave II** | *Slow* | 2.55 (0.14) | 2.60 (0.24) | 2.47 (0.18) | 2.57 (0.13) |
|  | *Fast* | 2.59 (0.22) | 2.62 (0.29) | 2.52 (0.21) | 2.64 (0.18) |
| **Wave III** | *Slow* | 3.64 (0.15) | 3.63 (0.22) | 3.57 (0.14) | 3.60 (0.16) |
|  | *Fast* | 3.82 (0.27) | 3.77 (0.28) | 3.70 (0.21) | 3.71 (0.17) |
| **Wave V** | *Slow* | 5.49 (0.21) | 5.37 (0.25) | 5.33 (0.22) | 5.37 (0.24) |
|  | *Fast* | 5.76 (0.29) | 5.70 (0.39) | 5.64 (0.27) | 5.64 (0.25) |
| **Wave V'** | *Slow* | 6.27 (0.24) | 6.14 (0.25) | 6.18 (0.27) | 6.22 (0.44) |
|  | *Fast* | 6.62 (0.37) | 6.50 (0.41) | 6.45 (0.34) | 6.47 (0.35) |
| **I–III IPI** | *Slow* | 2.26 (0.14) | 2.21 (0.16) | 2.23 (0.19) | 2.20 (0.15) |
|  | *Fast* | 2.42 (0.25) | 2.34 (0.27) | 2.31 (0.20) | 2.27 (0.17) |
| **III–V IPI** | *Slow* | 1.85 (0.14) | 1.74 (0.13) | 1.76 (0.17) | 1.77 (0.14) |
|  | *Fast* | 1.94 (0.23) | 1.93 (0.23) | 1.94 (0.23) | 1.93 (0.13) |
| **I–V IPI** | *Slow* | 4.11 (0.20) | 3.94 (0.20) | 3.99 (0.27) | 3.97 (0.22) |
|  | *Fast* | 4.36 (0.29) | 4.27 (0.29) | 4.25 (0.23) | 4.20 (0.23) |

Values are Mean (SD) in milliseconds. Slow rate = 19.1/s; Fast rate = 59.1/s. IPI = interpeak interval. Children ≤ 17 years; Adults ≥ 18 years.

**Figure 2:**
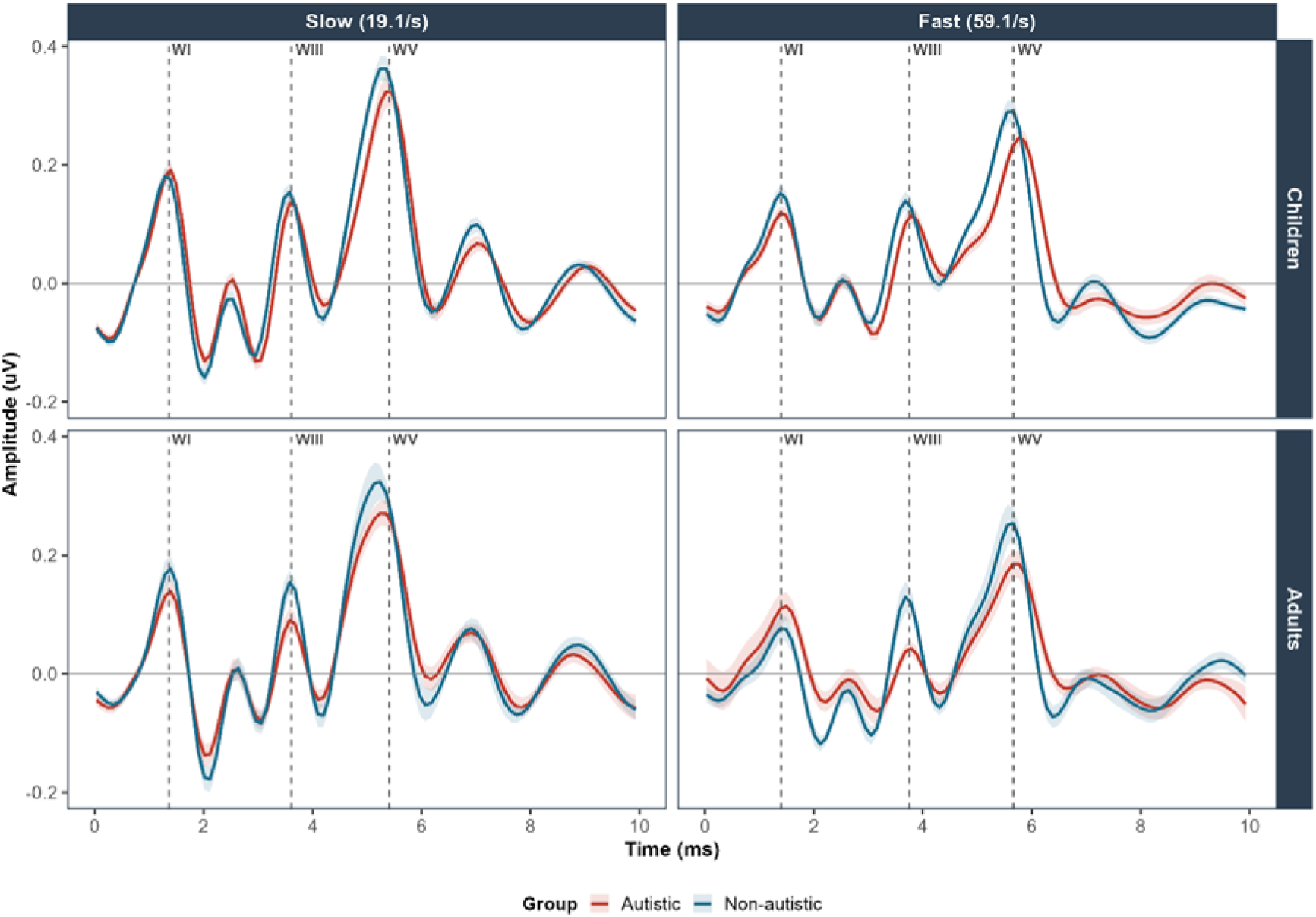
ABR Grand Average Waveforms by Group and Age Group. Slow rate = 19.1/s; Fast rate = 59.1/s | Shaded region = +/- 1 SE

Regarding absolute latencies, there was group x age interaction at Wave I (Figure 3; F (1, 125.20) = 6.37, p = .013, FDR p = .039) and group x age x rate at Wave V (Figure 3; F (1, 138) = 4.96, p = .028, p_adj_ = .083). Post hoc slope comparisons of Wave I revealed that non-Autistic participants showed a significant positive developmental trajectory at both fast rate (β = 3.77, SE = 1.25, CI[1.31, 6.23], t(159) = −2.52, p = .013) and slow rate (β = 2.86, SE = 1.24, CI[.40, 5.32], t(159) = −2.22, p = .028). Autistic participants showed flat trajectories at both rates (fast: β = −0.37, CI [−2.49, 1.75]; slow: β = −0.79, CI [−2.91, 1.33]). For absolute wave V at slow rate (t(186) = −1.80, p = .073), Autistic people had a significant negative slope (β = -2.17, SE = 1.03, CI[-4.20, -0.13]) and non-Autistic people had a positive slope (β = 0.72, SE = 1.22, CI[-1.69, 3.12]). A group × rate × sex interaction was observed at Wave I (Figure 4; F (1, 122.33) = 6.69, p = .011, FDR p = .033). *Post hoc* simple effects indicated that the group difference was specific to females at slow rate (t(159) = 2.47, p = .015, d = 1.69, 95% CI [0.33, 3.05]), with no group difference in males at either rate or in females at fast rate.

**Figure 3:**
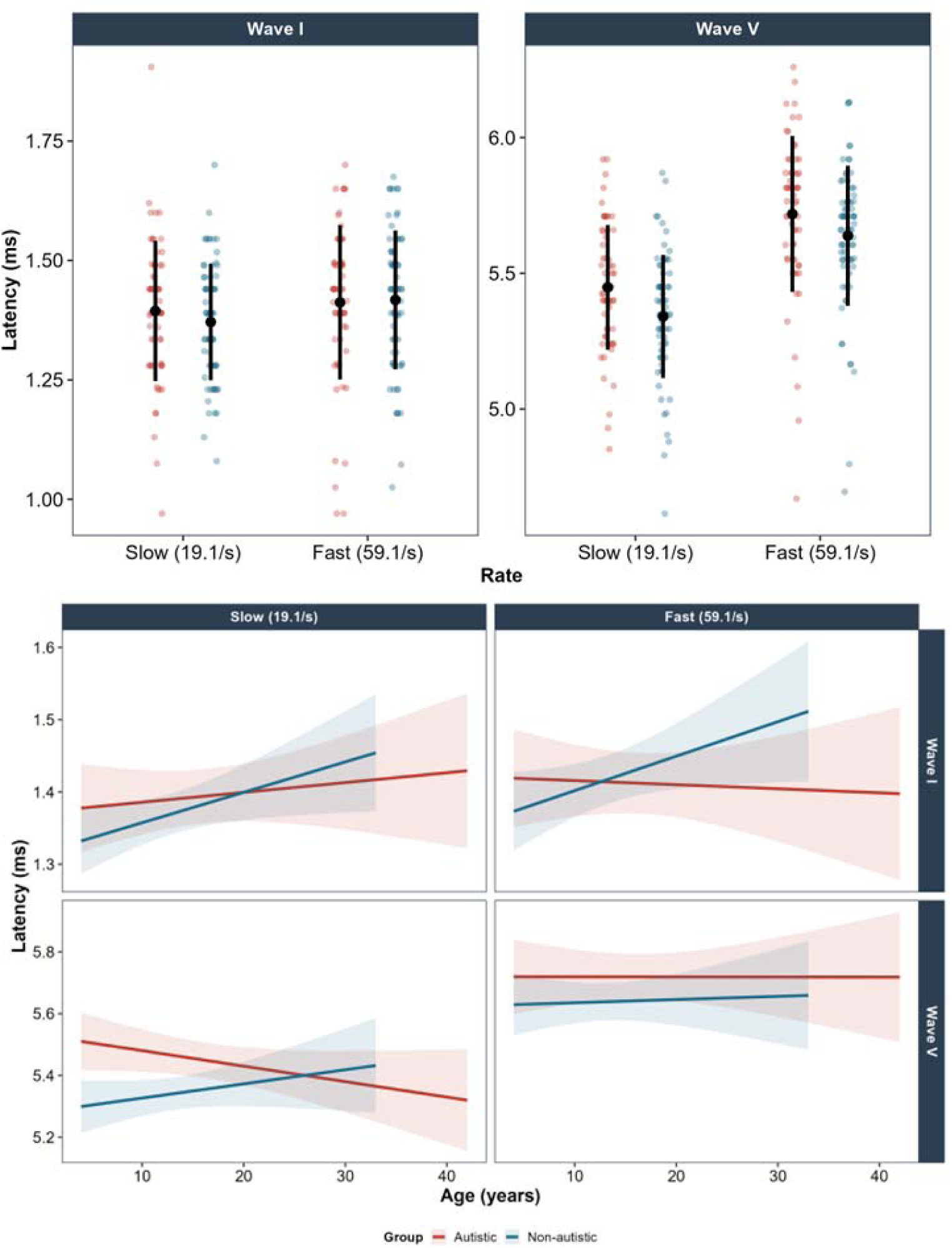
Wave I and V Latency Developmental Trajectories by Group and Rate. Top: Individual points with means ± SD Bottom: Regression lines ± 95% CI. Wave I: Group × Age F(1,125.20) = 6.37, p = .013 | Wave V: Group × rate × Age F(1,138) = 4.96, p = .028

In terms of interpeak latencies there was several interactions but no simple effects (Figure 5). For IPL I-III the was group x rate x sex (F(1, 137.23) = 4.41, p = .038, p_adj_ = 0.11). For IPL III-V group × rate (F(1, 138) = 5.35, p = .022, p_adj_ = .07) and a trending group × rate × age (F(1, 138) = 3.26, p = .073). An exploratory analysis of this interaction revealed at the slow rate a significant negative slope (β = -3.36, SE = 1.11, CI[-5.56, -1.17]) for the Autistic group while, non-Autistic group had non-significant flat slope (β = -.16, SE = 1.31, CI[-2.74, 2.43], t(222) = −1.86, p = .064). For IPL I-V there was a group × rate x age (F(1, 134.51) = 4.97, p = .028, p_adj_ = 0.08) interaction with no simple effects.

**Figure 4:**
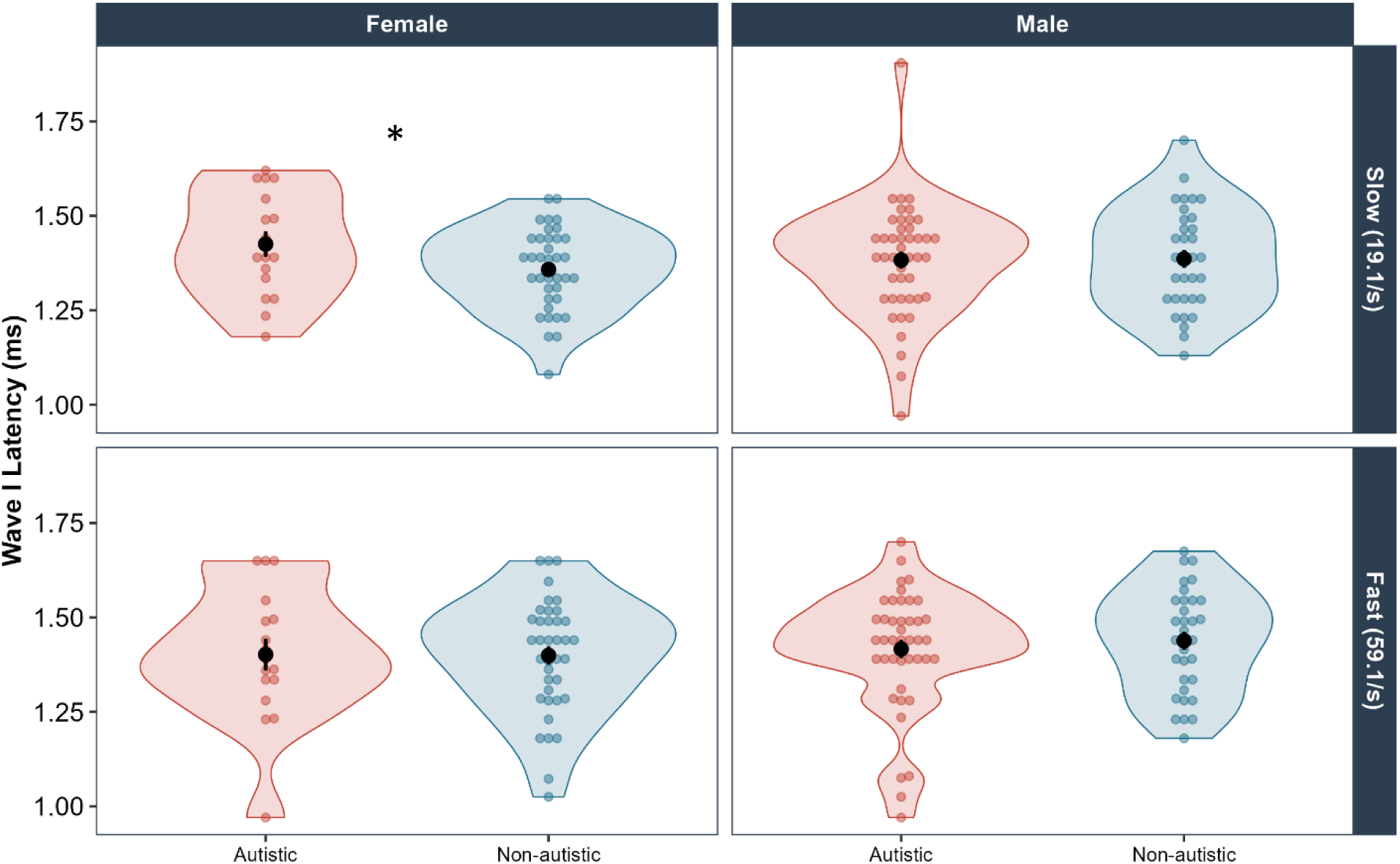
Wave I Latency by Group, Sex, and Rate. Group x rate x Sex: F(1,122.33) = 6.69, p = .011 | Female slow rate: t(159) = 2.47, p = .015, d = 1.69 | Individual points with means ± SE

**Figure 5:**
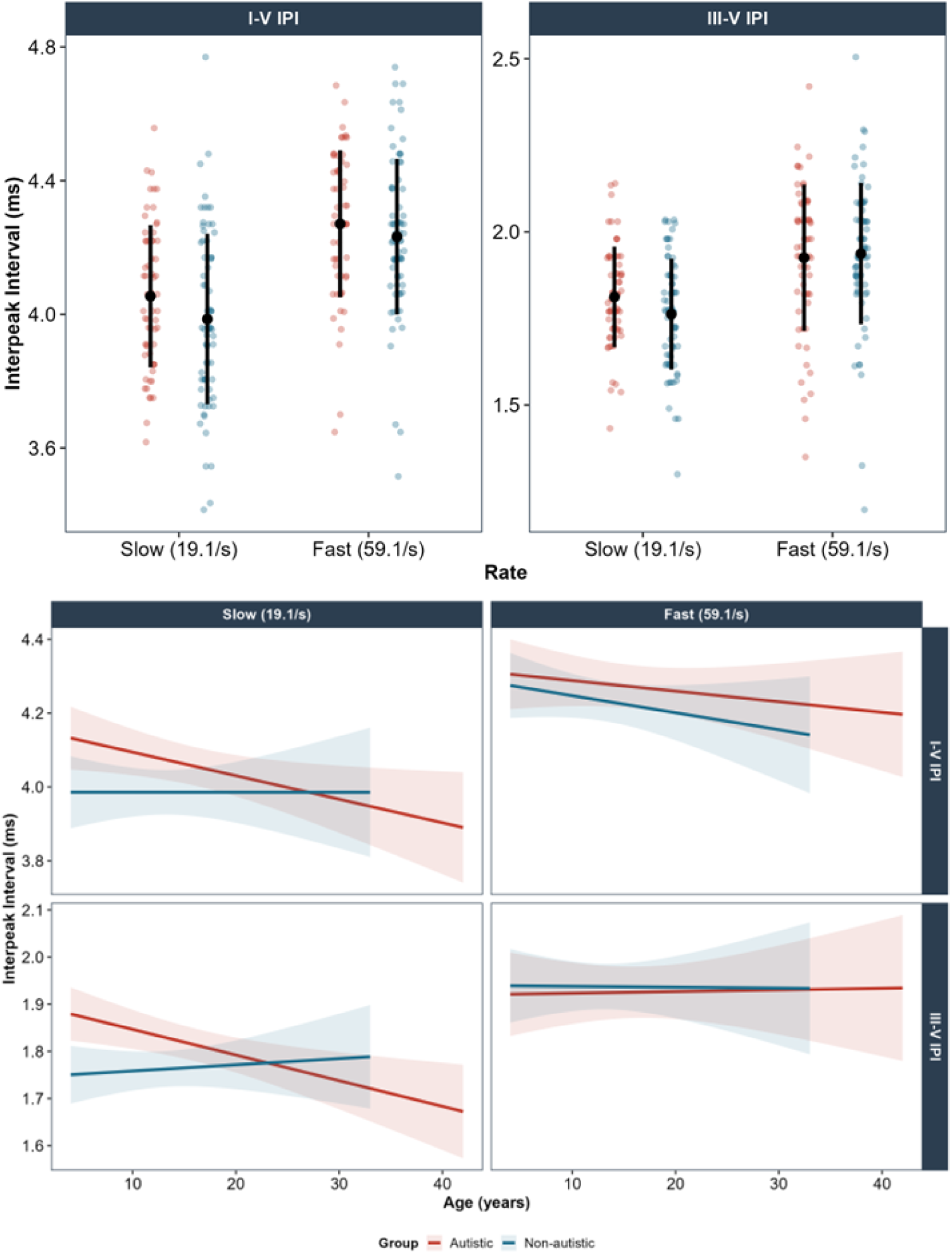
III-V and I-V Interpeak Interval Developmental Trajectories. Top: Individual points with means ± SD Bottom: Regression lines ± 95% CI.III-V IPI: Group x rate x Age p = .073 | I-V IPI: Group x rate x Age F(1,134.51) = 4.97, p = .028

Taken together, these findings suggest a pattern of atypical auditory brainstem development in autism characterised by diverging developmental trajectories at slow rate, with Autistic participants showing flatter or negative age-related changes in absolute and interpeak latencies relative to the positive trajectories observed in non-Autistic participants, a pattern that was most pronounced in females and specific to the slow stimulation rate, indicative of differences in neural synchrony under low-demand conditions.

### ABR: Wave amplitudes

The amplitudes of waves I, III, and V (sex × click-rate × diagnostic group × Age) were analyzed. There was a significant sex × click-rate × diagnostic group × Age interaction (F(1, 137.41) = 3.96, p = .048, p_adj_ = .15) for wave III amplitudes but no simple effects. There were no differences in amplitude of Waves I and V (Table 3).

**Table 3.** ABR Amplitude — Group Means (SD)

| Feature | Rate | Autistic |  | Non-Autistic |  |
| --- | --- | --- | --- | --- | --- |
|  |  | Children | Adults | Children | Adults |
| **Wave I** | *Slow* | 0.40 (0.15) | 0.35 (0.14) | 0.44 (0.18) | 0.41 (0.17) |
|  | *Fast* | 0.24 (0.09) | 0.20 (0.10) | 0.27 (0.11) | 0.25 (0.09) |
| **Wave III** | *Slow* | 0.28 (0.13) | 0.21 (0.10) | 0.35 (0.14) | 0.31 (0.14) |
|  | *Fast* | 0.20 (0.10) | 0.21 (0.18) | 0.23 (0.10) | 0.23 (0.11) |
| **Wave V** | *Slow* | 0.50 (0.25) | 0.39 (0.14) | 0.55 (0.18) | 0.47 (0.20) |
|  | *Fast* | 0.41 (0.19) | 0.30 (0.12) | 0.44 (0.15) | 0.38 (0.14) |
| **V/I ratio** | *Slow* | 1.35 (0.65) | 1.39 (0.76) | 1.52 (0.94) | 1.57 (1.36) |
|  | *Fast* | 2.57 (2.14) | 3.53 (5.51) | 2.26 (2.48) | 2.04 (1.77) |

Values are Mean (SD) in $\mu\text{V}$ . Slow rate = 19.1/s; Fast rate = 59.1/s. V/I ratio is dimensionless. Children $\leq 17$ years; Adults $\geq 18$ years.

### ASR: Startle Magnitude

Startle magnitude was higher for Autistic participants with a significant Group × Age interaction (F(1, 99) = 4.68, p = .033, η^2^p = .05; Figure 6) on acoustic reactivity. Autistic participants showed a positive age-related increase in startle amplitude (slope = +0.018 log units/year, SE = 0.011) while non-Autistic participants showed a negative age-related decrease (slope = −0.020 log units/year, SE = 0.014), with a significant between-group difference in developmental slope (estimate = 0.038, SE = 0.018, t(99) = 2.16, p = .033). At 105 dB, geometric means were 507 µV for Autistic adults versus 258 µV for non-Autistic adults (arithmetic means: 711 ± 690 µV vs 411 ± 560 µV), compared to 316 µV and 289 µV respectively in children (arithmetic means: 402 ± 301 µV vs 418 ± 478 µV), consistent with a group difference emerging in adulthood. There were no main effects of Group, Age, or Sex (all p > .32).

**Figure 6:**
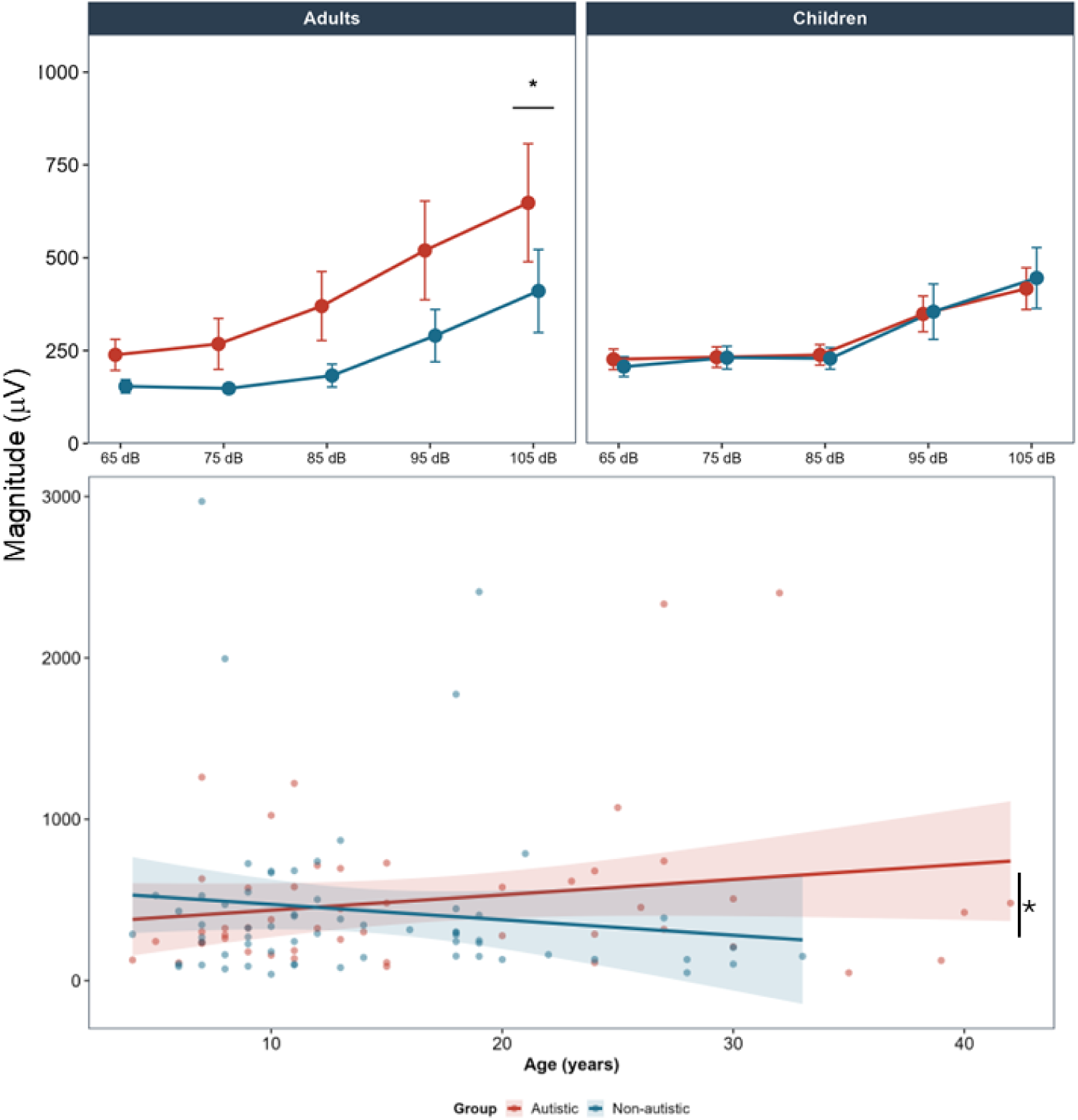
ASR amplitude across sound pressure levels and age by group. Top: Mean (± SE) startle magnitude across sound pressure levels (65–105 dB) for Autistic and non-Autistic participants, split by age group. Autistic adults showed significantly larger amplitude than non-Autistic adults at 105 dB (* p < .05, planned contrast). No group difference was observed in children. Bottom: Startle magnitude at 105 dB plotted against continuous age. Regression lines (± 95% CI) show diverging developmental trajectories: Autistic participants showed a positive age-related increase in amplitude (+0.018 log units/year) while non-Autistic participants showed a negative age-related decrease (−0.020 log units/year); the between-group difference in slope was significant (* p = .033). Mean ± SE shading. Children: n = 42/54; Adults: n = 17/24 (Autistic/Non-Autistic).

### ABR and acoustic reactivity associations

ABR amplitudes and acoustic reactivity associations are in opposite directions for Autistic and non-Autistic participants. Wave I and Wave V amplitudes were positively correlated with reactivity for the non-Autistic participants (Figure 7; Wave I: rho = +0.300, p = .026; Wave V: rho = +0.253, p = .063) while they were negatively correlated with reactivity for the Autistic participants (Wave I: rho = −0.193, p = .194; Wave V: rho = −0.204, p = .169). The associations were significantly different from each other (Wave I: z = −2.462, p = .014, p_adj_ = .035; Wave V: z = −2.271, p = .023, p_adj_ = .035).

**Figure 7:**
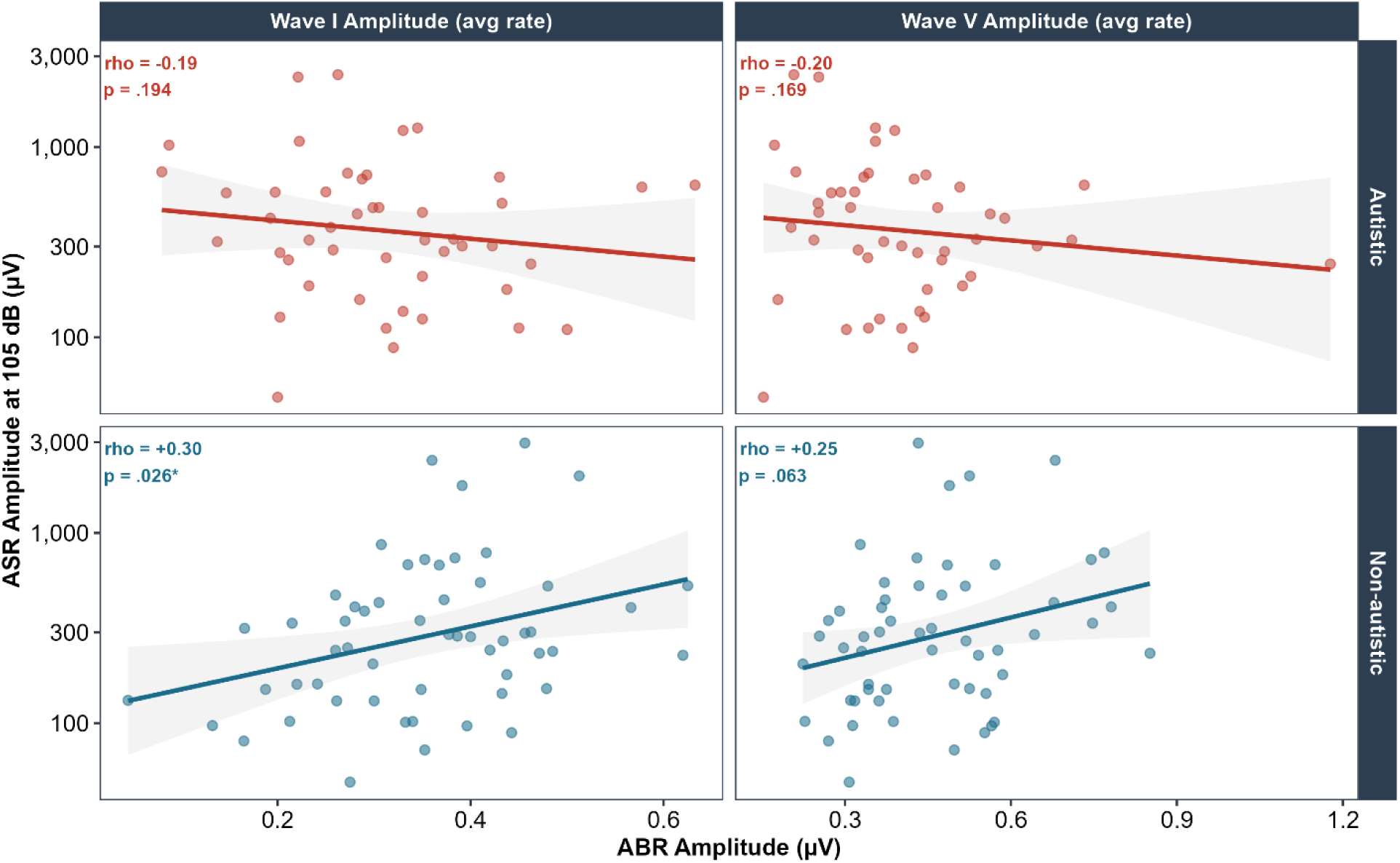
ABR Amplitude μV ASR Amplitude — Opposite Associations by Group. Fisher’s z: Wave I z = -2.46, p = .014 | Wave V z = -2.27, p = .023 | Regression lines (± 95% CI)

## Discussion

We investigated whether Autistic children and adults showed a combination of brainstem-mediated auditory timing differences and heightened behavioral reactivity to sound. Participants completed a passive auditory paradigm including ABR as a measure of brainstem neural timing and synchrony, and ASR as a measure of behavioral reactivity. We hypothesized that Autistic children would present with altered brainstem auditory processing while Autistic adults would show normalized brainstem responses alongside elevated behavioral reactivity. This profile would be consistent with compensatory adaptive auditory gain formed during critical windows of experience-induced plasticity in early development, which persists and becomes maladaptive in adulthood as brainstem responses normalize. The auditory system expresses increased experience-induced plasticity in younger age. For example, in congenitally deaf children, the maturation of the auditory evoked potentials depended on the age of cochlear implantation with optimal development occurring in implantation before the age of 3.5 years^75^. The findings partially support this hypothesis, with some important distinctions from our original predictions.

Autistic participants showed differences in auditory brainstem timing relative to non-Autistic participants, though some of these did not survive FDR correction after multiple comparison adjustment. Exploratory *post hoc* analyses revealed some rate-specific and sex-specific patterns. There were no group differences in ABR amplitudes.

The most consistent pattern across the ascending brainstem pathway was a divergence in developmental trajectories at slow rate. Non-Autistic participants showed positive age-related increases in Wave I latency at both click rates, a trajectory consistent with known increases in early ABR components across development^76^. However, Autistic participants showed flat trajectories at both rates, with the group contrast reaching significance at both slow and fast rate. The persistence of wave I differences across development aligns with the literature in which atypical wave I latencies in ASD children of all ages remains the most common outcome^77^. Similarly, at the level of the inferior colliculus (Wave V) and the interpeak intervals spanning the central brainstem pathway (III–V IPI, I–V IPI), Autistic participants showed significantly negative developmental slopes at slow rate, while non-Autistic participants showed flat or positive trajectories. These effects were absent at fast rate.

The interpretation of these developmental trajectory differences is not straightforward. A divergence in trajectories does not imply a simple group difference rather it implies that the two groups are on different maturational paths that are moving apart rather than converging. It is important to note that the majority of ABR maturation occurs in the first three to five years of life^78^, driven by rapid myelination and auditory pathway refinement, meaning that by the time participants enter our sample at age four, the majority of maturational changes are largely complete. The trajectory differences observed here therefore reflect more gradual, later-stage developmental and aging-related changes rather than early maturation. In non-Autistic participants, the positive age-related increase in Wave I latency across the sample range is consistent with known effects of aging on peripheral auditory nerve conduction, in which latency increases progressively from childhood through adulthood as the cochlea and auditory nerve undergo age-related changes^76,79^. The stability of Wave V latency in non-Autistic participants aligns with previous reports^76^. In Autistic participants, the flat or negative trajectories across this same age range suggest a divergence from this normative aging pattern and not necessarily a failure of early maturation, but a different relationship between age and brainstem timing that becomes apparent when examined continuously across the developmental span. Whether this reflects a genuinely different developmental trajectory, a ceiling effect from atypical early maturation that was already present before age four, or a combination of both, cannot be determined from the current cross-sectional data. This is also broadly consistent with studies reporting a bigger group difference in participant samples younger than eight ^66^, and reinforces the need for studies beginning in infancy to capture the full developmental arc.

Another important finding was the female-specific group difference in Wave I latency at slow rate. Autistic females showed significantly longer Wave I latency than non-Autistic females at slow rate, while no group difference was observed in males at either rate. This female-specific pattern was consistent across Wave I and Wave V which suggests a consistent pattern. The Autistic female sample was small, and this finding requires replication. It is important to investigate this difference given known sex differences in ABR morphology, with females typically showing shorter latencies^80,81^ and larger amplitudes than males^81,82^, and evidence that the female Autistic phenotype may be neurobiologically distinct from that of Autistic males^83,84^.

The trending nature of the ABR findings of effects that are directionally consistent and, in some cases, show large effect sizes, particularly in females, but do not survive multiple comparison correction is likely partly a power problem rather than an absence of a true difference. The Autistic group in this study was heterogeneous in both age and phenotype, spanning a wide developmental range and including participants with varying degrees of sensory and cognitive profiles. Power to detect group differences in ABR features is not simply a function of total sample size but of the homogeneity of the signal within the Autistic group. A larger sample of Autistic participants with a more defined sensory profile such as those with documented auditory hypersensitivity or sensory over-responsivity would likely show more robust and consistent ABR differences, as the signal would not be diluted by participants whose auditory processing profile is closer to the non-Autistic range. Sensory phenotyping studies report subgroups that experience high sensory sensitivities such as generalized sensory differences subgroup and under-responsive and sensation seeking subgroup, in contrast to a sensory adaptive subgroup that has sensory phenotype closer to non-Autistic people^85^. The differences in the sensory phenotypes are also linked to differences in functional connectivity across cortical, subcortical, and network levels^86^. This is a known challenge in autism research more broadly: heterogeneity within the diagnostic category attenuates group-level effects and inflates the sample size required to detect them. The trending effects reported here may therefore represent genuine neurobiological differences that would reach conventional significance thresholds in a larger, more phenotypically homogeneous Autistic sample.

In terms of the ASR, ASR amplitude showed diverging developmental trajectories between groups, with Autistic participants showing a positive age-related increase in startle magnitude while non-Autistic participants showed a negative age-related decrease. No group difference was observed in children, and the divergence emerged continuously across development rather than appearing discretely in adulthood. ASR latency was equivalent across groups and ages, confirming that Autistic participants are not slower to initiate their startle response. They startle more intensely with age, but at the same speed.

The developmental trajectory of the ASR group difference is central to the adaptive gain interpretation. If hyperreactivity were a fixed trait, a group difference would be expected uniformly across age. Instead, the groups show overlapping amplitude in childhood and diverge progressively across development, at the same developmental period during which ABR timing trajectories show divergence between groups. This temporal co-occurrence is consistent with the hypothesis that compensatory gain upregulation, initially adaptive in response to early brainstem timing differences, becomes maladaptive as the brainstem partially normalizes across development.

The ABR and ASR pathways directly intersect at the cochlear nerve and the cochlear nucleus. They also indirectly intersect through the midbrain ASR modulatory pathways (inferior colliculus - superior colliculus - Pedunculopontine Tegmental Nucleus) that adjust the response intensity based on the sensory context. The increased magnitude of the startle response associated with Autism is regulated by the excitability, size and number of recruited PnC giant neurons, as recently shown in an *Cntnap2* knock-out Autism rat model^92^. A similar modulation in the sensorimotor interface of those pathways is most likely causing the increased reactivity to sound in Autistic individuals.

The relationship between ABR amplitude and ASR amplitude showed opposite directions in Autistic and Non-Autistic participants, and this group difference survived FDR correction. In non-Autistic participants, larger brainstem amplitude was positively associated with larger startle reactivity at both Wave I and Wave V, with the between-group difference in correlation direction being FDR-significant for both waves. In Autistic participants, the same correlations were negative, indicating that those with larger brainstem responses showed smaller startle reactivity which is counterintuitive.

In non-Autistic participants, the positive relationship between ABR and ASR amplitude makes intuitive sense as someone with a stronger brainstem response to sound also startles more strongly, which is what you would expect if individual differences in auditory sensitivity are consistent across measures. The reversal of this pattern in Autistic participants is harder to explain. Rather than a stronger brainstem response predicting a stronger startle, the relationship goes the other way in which those with larger brainstem responses tend to show smaller or equivalent startle magnitude. This suggests that in Autism, the size of the brainstem response does not determine how strongly the startle circuit responds downstream. Whatever is driving the elevated startle reactivity in Autistic adults appears to operate separately from, or in opposition to, the magnitude of the incoming brainstem signal which points toward a difference in how gain is regulated at the central level rather than a simple amplification of peripheral input.

The idea of maladaptive sensory gain stems from reports of reduced neurotransmission of the ascending auditory pathway of newborn babies later diagnosed with Autism^58,93^. During early postnatal development neural circuits, both structural and functional, are shaped by exposure to sensory input. These periods of heightened plasticity play a crucial role in the maturation of sensory representations in both subcortical and cortical regions, influencing the development of behavior and cognition. In the auditory system, this period of rapid refinement supports the acquisition of specialized skills such as phoneme recognition, language learning, absolute pitch, and musical ability, but it also marks a time of heightened vulnerability to sensory deprivation. The early reduced responsivity and slowed neurotransmission regardless of the molecular reason could induce adaptive excitatory gain or reduced inhibitory gain within the auditory system. The ABR pathway partially recovers its output and therefore what was considered adaptive would induce maladaptive gain, leading to behavioral hyper-reactivity.

Mechanistically, the ABR differences could be linked to possible molecular and cellular changes. Slower neural transmission could be due to dysfunction in myelination, pathway length, axonal diameter, the (a)synchronization of neuronal firing, or synaptic efficacy^94^. Those factors cannot be directly linked or consistently linked to Autism as there is evidence of opposing findings within the central nervous system. For example, diffusion tensor imaging mean diffusivity of white matter microstructure has been reported as both increased^95^ and decreased^96^ in Autism. While ABRs do not directly measure white matter microstructure, ABRs are sensitive to changes in axonal morphology and myelination, axon bundle density and fiber orientation distribution, and other intra- and extra-cellular processes.

There is no consensus on the structural or synaptic condition within the ascending brainstem pathways in Autism therefore it is difficult to disentangle the specific reasons for the reduction in action potential velocity. One interpretation of our findings is that the ascending auditory pathway has a combination of abnormalities such as reduced number of cellular nuclei, reduced axonal diameter, asynchronization of neuronal firing and changes in myelination that results in this unique ABR profile in Autism. Those kinds of changes are observed in post-mortem analysis of the superior olivary complex in which the nuclei have altered volumes^44,98^, shapes^45^, and number^45^ (for a review^59^). Some of those features undergo tuning based on sensory experiences and this experience-based tuning may lead to cascading sensory and hyperreactivity differences in Autistic children and adults.

### Addressing ABR Methodology Heterogeneity

The ABR literature in Autism is marked by substantial inconsistency, with studies reporting increased, decreased, or absent latency and amplitude differences across samples and methodologies^59,66^. Age range, diagnostic criteria, and peripheral hearing screening have been identified as contributing factors^59,66^, but one underappreciated source of variation is the stimulus click rate. Click rate directly determines the degree of neural challenge imposed on the auditory nerve: faster rates require more rapid neural recovery and temporal following, producing longer latencies and smaller amplitudes. Studies using slow rates, fast rates, or a single rate without comparison are not directly comparable, and pooling findings across rate conditions obscures real rate-dependent effects.

This study employed both a slow (19.1/s) and fast (59.1/s) click rate within the same participants, allowing rate-dependent effects to be distinguished from rate-independent ones. Click rate had a significant effect on ABR features, which is consistent with what is already known about rate-dependent neural slowing^90^. More importantly, the group differences in developmental trajectories were specific to the slow rate. Autistic participants showed flat or negative age-related slopes in Wave I latency and interpeak intervals at slow rate, while these patterns were absent at fast rate. This rate specificity would be missed in single-rate designs and may explain why studies using only fast rates report less consistent group differences, while slow-rate studies are more likely to detect timing-related effects.

Beyond the methodological point, the slow-rate specificity tells us something about the nature of the difference itself. Slow-rate stimulation is more sensitive to baseline neural synchrony and conduction, whereas fast-rate stimulation reflects capacity under increased temporal demand^91^. Given that group differences emerge primarily at slow rate suggests that the Autistic auditory brainstem differs in its baseline temporal processing rather than in its response to neural load. This has direct implications for ABR protocol design in future autism research in which a single fast-rate condition is likely to enough to capture developmentally relevant group differences, and inclusion of a slow-rate condition should be considered standard.

### Limitations

This study has several limitations that warrant consideration when interpreting the findings. First, the exclusion of Autistic participants with the most severe auditory and sensory sensitivities introduces a potential selection bias. Some participants, particularly those with significant tactile sensitivities, were unable to complete the study due to discomfort caused by the conductive gel, despite the study being focused on auditory processing. This limits the generalizability of the results, as the participants who completed the study may represent a subset of the Autistic population with milder sensory challenges. In addition, the Autistic female subsample was small, meaning the female-specific latency effects, while striking in magnitude, are based on cell sizes that preclude definitive conclusions. Moreover, the cross-sectional design means that developmental convergence is inferred from age-group differences rather than individual change over time, longitudinal data would be required to confirm that individual Autistic participants’ brainstem timing trajectories approach non-Autistic values across development. Finally, the ABR–ASR correlation analyses are cross-sectional and correlational; therefore, no causal inference about the relationship between brainstem processing and startle gain is possible.

## Conclusion

This study provides preliminary evidence for atypical auditory brainstem timing in Autism, characterised by diverging developmental trajectories at slow stimulation rate and a female-specific group difference in early auditory nerve latency. Importantly, these timing differences occur in the absence of any group differences in brainstem response magnitude, dissociating temporal and magnitude-based aspects of auditory brainstem processing in Autism. In adulthood, Autistic participants show significantly elevated startle reactivity despite equivalent ASR timing, consistent with the hypothesis that compensatory central gain in which upregulation in response to early brainstem timing differences becomes maladaptive as brainstem timing normalises across development. While the findings require replication, they collectively support a developmental model in which early atypicalities in auditory brainstem timing have cascading effects on sensory reactivity that persist and amplify into adulthood.

## Financial Disclosures / Acknowledgements

RAS is funded through a Dorothy Killam Fellowship, an NSERC Discovery Grant (RGPIN-2024-06233), two SSHRC Insight Grants (435-2017-0936 & 435-2024-1375), a CIHR Project Grant (487850), the University of Western Ontario Faculty Development Research Fund, and a Canadian Foundation for Innovation John R. Evans Leaders Fund (37497), and through a grant from the Canada First Research Excellence Fund (BrainsCAN). RS would also like to thank M Rankin, K MacLellan, A O’Hanley, and S Riley for Alvvays supporting us.^[^. The study was also supported by a CIHR project grant to SS (168866) and the Simons Foundation for Autism Research (SFARI). The remaining authors reported no biomedical financial interests or potential conflicts of interests.

## CRediT Statement

AS – Conceptualization, Methodology, Formal analysis, Investigation, writing – original draft, writing – review and editing, Visualization. RG – Investigation, MR – Investigation, CM – Investigation, KS – Investigation, SES – Conceptualization, methodology, and being our guardian angel. RAS – Conceptualization, resources, writing – review and editing, supervision, project administration, funding acquisition. SS – Conceptualization, review and editing

